# Circulating Mitochondrial DNA is an Early Indicator of Severe Illness and Mortality from COVID-19

**DOI:** 10.1101/2020.07.30.227553

**Authors:** Davide Scozzi, Marlene Cano, Lina Ma, Dequan Zhou, Ji Hong Zhu, Jane A O’Halloran, Charles Goss, Adriana M. Rauseo, Zhiyi Liu, Valentina Peritore, Monica Rocco, Alberto Ricci, Rachele Amodeo, Laura Aimati, Mohsen Ibrahim, Ramsey Hachem, Daniel Kreisel, Philip A. Mudd, Hrishikesh S. Kulkarni, Andrew E. Gelman

## Abstract

Mitochondrial DNA (MT-DNA) are intrinsically inflammatory nucleic acids released by damaged solid organs. Whether the appearance of cell-free MT-DNA is linked to poor COVID-19 outcomes remains undetermined. Here, we quantified circulating MT-DNA in prospectively collected, cell-free plasma samples from 97 subjects with COVID-19 at the time of hospital presentation. Circulating MT-DNA were sharply elevated in patients who eventually died, required ICU admission or intubation. Multivariate regression analysis revealed that high circulating MT-DNA levels is an independent risk factor for all of these outcomes after adjusting for age, sex and comorbidities. Additionally, we found that circulating MT-DNA has a similar or superior area-under-the curve when compared to clinically established measures of systemic inflammation, as well as emerging markers currently of interest as investigational targets for COVID-19 therapy. These results show that high circulating MT-DNA levels is a potential indicator for poor COVID-19 outcomes.

## INTRODUCTION

Coronavirus Disease 2019 (COVID-19) is a respiratory tract infection caused by severe acute respiratory syndrome coronavirus 2 (SARS-CoV-2) that has resulted in a global health emergency, causing considerable strain on economic, social and medical systems (Lipman et al., 2020). COVID-19 presents in a wide spectrum of severity. Although most patients develop only mild or uncomplicated illness, others require prolonged hospitalization, ICU care and intubation for respiratory support. In severe cases, patients can develop acute respiratory distress syndrome (ARDS), cytokine storm, multi-organ failure and death (Lipman et al., 2020). Although the underlying mechanisms of severe COVID-19 illness remain unclear, it appears to be exacerbated by an over-exuberant innate immune response (Vabret et al., 2020). These observations have led to several ongoing clinical trials targeting components of the innate immune response such as inflammatory cytokine signaling and complement activation (Maes et al., 2020; Rilinger et al., 2020; Smith et al., 2020).

Previous work has established that viral infection can trigger cellular necrosis, which in turn inhibits viral replication along with amplifying anti-viral immune responses through the release of damage associated molecular patterns (DAMPs) (Nailwal and Chan, 2019). DAMPs in particular are potent triggers of innate responses through their engagement of pattern recognition receptors such as toll-like receptors (TLR) that drive the expression of inflammatory cytokines and presentation of viral antigens (Kulkarni et al., 2020a). Mitochondrial DNA (MT-DNA) is a member of a group of mitochondrial DAMPs (MT-DAMPs) released by injured or dying cells and is recognized by TLR9 due to encoded hypomethylated CpG motifs reminiscent of an ancestral bacterial origin (Zhang et al., 2010). MT-DNA levels have been previously shown to be elevated in the plasma of patients that develop ARDS and multi-organ dysfunction during sepsis, as well as during sterile injury including trauma, hemorrhagic shock and ischemia-reperfusion (Faust et al., 2020; Hauser et al., 2010; Nakahira et al., 2013; Puskarich et al., 2012; Scozzi et al., 2019; Simmons et al., 2013). The release of MT-DNA is often accompanied by the release of other MT-DAMPs, such as N-formylated peptides, cytochrome c and cardiolipin, which collectively engage multiple TLRs and the N-formylated peptide receptor-1 that in turn not only induce inflammatory cytokine expression (Grazioli and Pugin, 2018; Wu et al., 2019; Zhang et al., 2014) but also the generation of reactive oxygen species and the facilitation of neutrophil trafficking and activation (Dorward et al., 2017; Nakayama and Otsu, 2018; Scozzi et al., 2019). Through these effects, MT-DAMPs can directly contribute to acute lung injury and systemic inflammation (Zhang et al., 2010). Given that ARDS secondary to SARS-CoV-2 infection is also linked with lung tissue injury and immune cell activation (Mangalmurti and Hunter, 2020), we asked if elevated levels of circulating MT-DNA could be used as a risk indicator for the development of severe illness.

Here, we demonstrate that COVID-19 patients with high circulating levels of MT-DNA are more likely to require ICU level care and intubation and are at heightened risk of death. In addition, MT-DNA quantitation as predictor of poor COVID-19 outcomes is comparable or better than inflammation indicators commonly used in current clinical practice, as well as certain emerging immune markers.

## RESULTS

### Participants and Descriptive Data

A total of 107 subjects were assessed for eligibility from 3/26/2020 to 4/26/2020. Of these, 97 adult subjects with laboratory-confirmed COVID-19 were included in our study (**Table 1, Figure 1**). 10 subjects were excluded as they did not have samples from the day of presentation. The median age of the population in our study was 65 (54-73). Among these subjects, 55.6% (54/97) were male, 77.3% (75/97) were African-American, 46.4% (45/97) were obese (BMI ≥ 30), and 46.3% (45/97) had a positive smoking history. The median follow up time was 81 days (74-87).

**Table 1.** Demographic and Clinical characteristics associated with mortality.

|  | **Total**<br>**(n=97)** | **Alive**<br>**(n=72)** | **Deceased**<br>**(n=25)** | **OR for mortality**<br>**(95% CI)** | **P**<br>**value** |
| --- | --- | --- | --- | --- | --- |
| **Demographics** |  |  |  |  |  |
| Age, yr<sup>a</sup> | 65 (54-73) | 61.1 (50.3-70.5) | 76.2 (65.3-84.9) | 1.08 (1.04-1.13) | 0.0003 |
| Male n (%)<sup>b</sup> | 54 (55.6) | 39 (54.17) | 15 (60) | 1.26 (0.5-3.27) | 0.6 |
| African-American<sup>b</sup> | 75 (77.3) | 57 (79.2) | 18 (72) | 0.6 (0.24-2) | 0.4 |
| BMI<sup>a</sup> | 29.1 (24.7-34.4) | 28.9 (24.7-33.3) | 30.9 (24.5-37.6) | 1.02 (0.96-1.08) | 0.4 |
| Smoking history<sup>b</sup> | 45 (46.3) | 29 (40.3) | 16 (64) | 2.63 (1.04-7) | 0.04 |
| **Comorbidities** |  |  |  |  |  |
| HTN<sup>b</sup> | 73 (75.2) | 51 (70.8) | 22 (88) | 3.02 (0.91-13.71) | 0.09 |
| DMII<sup>b</sup> | 51 (52.5) | 33 (45.8) | 18 (72) | 3.04 (1.17-8.64) | 0.02 |
| COPD<sup>b</sup> | 13 (14.4) | 7 (9.7) | 6 (24) | 2.93 (0.85-9.89) | 0.08 |
| CKD>2<sup>b</sup> | 37 (38.1) | 24 (33.3) | 13 (52) | 2.16 (0.85-5.53) | 0.1 |
| ESRD<sup>b</sup> | 8 (8.2) | 4 (5.6) | 4 (16) | 3.23 (0.71-14.80) | 0.1 |
| CAD<sup>b</sup> | 26 (26.8) | 14 (19.4) | 12 (48) | 3.8 (1.44 -10.34) | 0.007 |
| CVA<sup>b</sup> | 21 (21.6) | 14 (19.4) | 7 (28) | 1.61 (0.54-4.53) | 0.3 |
| VTE<sup>b</sup> | 16 (16.4) | 14 (19.4) | 2 (8) | 0.36 (0.05-1.42) | 0.1 |
| 2 or more<br>comorbidities<sup>b</sup> | 62 (63.9) | 41 (56.9) | 21 (84) | 5.54 (1.72-24.91) | 0.009 |
| 3 or more<br>comorbidities<sup>b</sup> | 50 (51.5) | 32 (43.8) | 18 (75) | 3.21 (1.23-9.15) | 0.02 |
| Definition of Abbreviations: HTN= Hypertension; DMII= Diabetes Mellitus Type II; COPD= Chronic Obstructive Pulmonary Disease; CKD>2= Chronic Kidney Disease; ESRD= End-Stage Renal Disease; CAD= Coronary Artery Disease; CVA= Cerebrovascular Accident; VTE= Venous Thromboembolism. Continuous variables are reported as median (interquartile range). |  |  |  |  |  |
| P values indicate differences between alive and deceased COVID 19 patients. |  |  |  |  |  |
| <sup>a</sup>Values presented as median and interquartile range |  |  |  |  |  |
| <sup>b</sup>Values presented as number and % of the column total |  |  |  |  |  |

**Figure 1.**
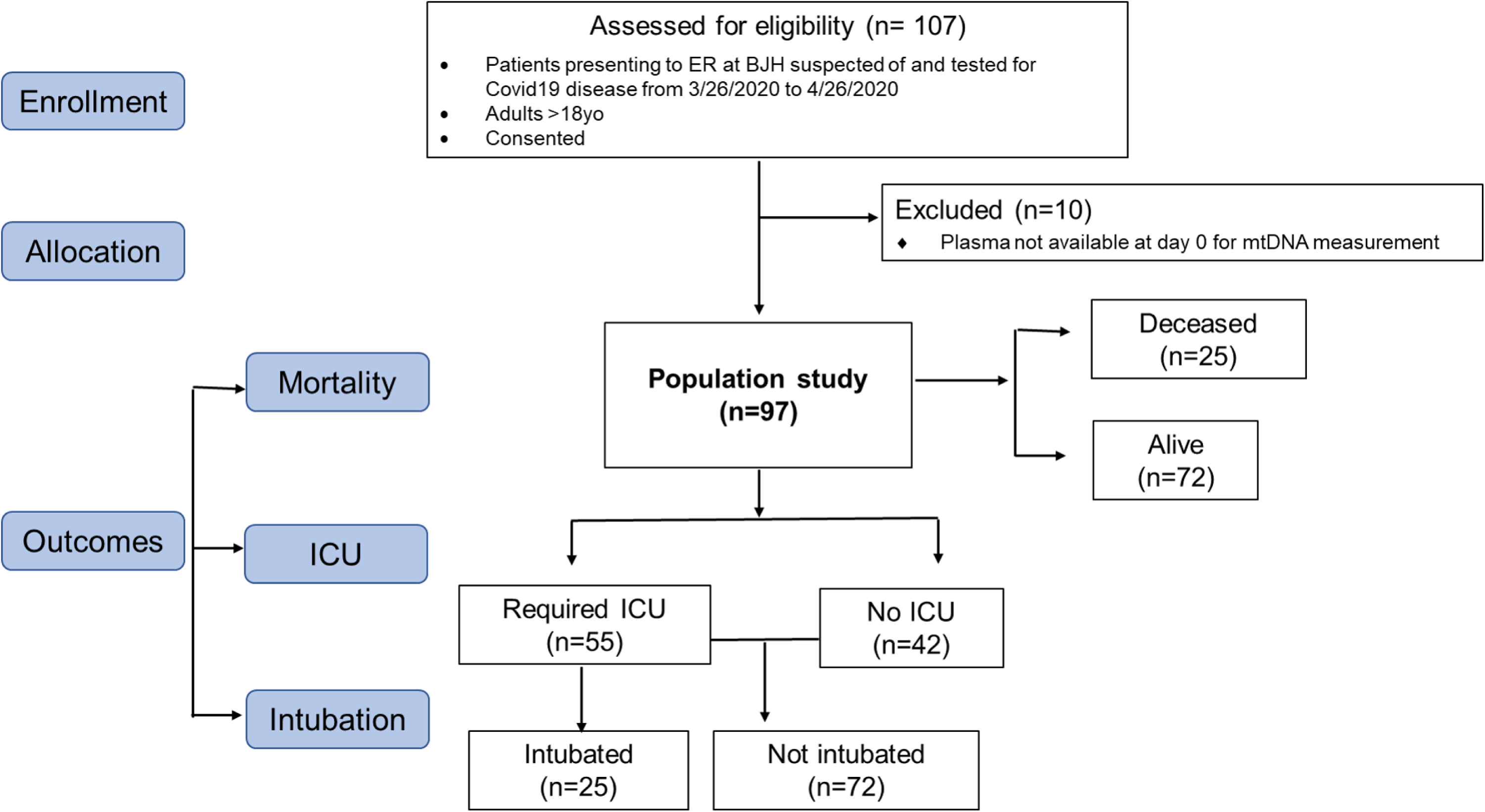
Consort flow diagram showing enrollment of patients, allocation, and outcomes.□.

### Outcome Data

The primary outcome of mortality was observed in 25.8% (25/97) subjects in our study (**Table 1**). Those who died were older [76.2 versus 61.1 years, OR 1.08 (1.04-1.13), p = 0.0003], were more likely to be smokers [64% versus 40.3%, OR 2.63 (1.04-7), p = 0.04] and have a higher proportion of type 2 diabetes mellitus [72% versus 45.8%, OR 3.04 (1.17-8.64), p = 0.02], coronary artery disease [48% versus 19.4%, OR 3.8 (1.44-10.34), p = 0.007], and 2 or more comorbidities [84% versus 56.9%, OR 5.54 (1.72-24.91, p = 0.009] on univariate analysis (**Table 1**).

56.7% of subjects with COVID-19 (55/97, **Table 2**) required an ICU admission. These subjects were older [71 versus 54.4, OR 1.1 (1.06-1.15), p <0.0001] and were more likely to have type 2 diabetes mellitus [65.5% versus 35.7%, OR 3.41 (1.49-8.07), p = 0.004], and 2 or more comorbidities [72.7% versus 52.4%, OR 2.66 (1.14-6.38), p = 0.02]. 25.8% (25/97) required invasive mechanical ventilation as a treatment for acute respiratory failure. The main clinical characteristic associated with higher risk for intubation was age, with a median of 70 years for intubated subjects compared with 61 years for non-intubated subjects [OR 1.05 (1.01-1.09), p = 0.005] on univariate analysis, **Table 3]**.

**Table 2.** Demographic and Clinical characteristics associated with ICU admission.

|  | No ICU<br>(n=42) | ICU<br>(n=55) | OR for ICU<br>(95% CI) | P value |
| --- | --- | --- | --- | --- |
| **Demographics** |  |  |  |  |
| Age, yr<sup>a</sup> | 54.4 (38.5-64.2) | 71 (63.8-78.9) | 1.1 (1.06-1.15) | <0.0001 |
| Male n (%)<sup>b</sup> | 19 (45.2) | 35 (63.6) | 2.11 (0.93-4.86) | 0.07 |
| African-American<sup>b</sup> | 33 (78.6) | 42 (76.4) | 0.88 (0.32-2.29) | 0.8 |
| BMI<sup>a</sup> | 30.2 (25.3-34.8) | 28.4 (24.4-33) | 0.98 (0.92-1.03) | 0.5 |
| Smoking history<sup>b</sup> | 15 (35.7) | 30 (54.5) | 2.16 (0.95-5) | 0.067 |
| **Comorbidities** |  |  |  |  |
| HTN<sup>b</sup> | 29 (69) | 44 (80) | 1.79 (0.7-4.61) | 0.2 |
| DMII<sup>b</sup> | 15 (35.7) | 36 (65.5) | 3.41 (1.49-8.07) | 0.004 |
| COPD<sup>b</sup> | 6 (14.3) | 7 (12.7) | 0.87 (0.26-2.93) | 0.8 |
| CKD>2<sup>b</sup> | 12 (28.6) | 25 (45.5) | 2.08 (0.89-5) | 0.09 |
| ESRD<sup>b</sup> | 4 (9.5) | 4 (7.3) | 0.74 (0.16-3.33) | 0.7 |
| CAD<sup>b</sup> | 8 (19.1) | 18 (32.7) | 2.06 (0.81-5.61) | 0.1 |
| CVA<sup>b</sup> | 7 (16.7) | 14 (25.5) | 1.7 (0.63-4.94) | 0.3 |
| VTE<sup>b</sup> | 8 (19.1) | 8 (14.6) | 0.72 (0.24-2.15) | 0.5 |
| 2 or more<br>comorbidities<sup>b</sup> | 22 (52.4) | 40 (72.7) | 2.66 (1.14-6.38) | 0.02 |
| 3 or more<br>comorbidities<sup>b</sup> | 16 (38.1) | 34 (61.8) | 2.63 (1.16-6.11) | 0.02 |
| Definition of Abbreviations: HTN= Hypertension; DMII= Diabetes Mellitus Type II; COPD= Chronic Obstructive Pulmonary Disease; CKD>2= Chronic Kidney Disease; ESRD= End-Stage Renal Disease; CAD= Coronary Artery Disease; CVA= Cerebrovascular Accident; VTE= Venous Thromboembolism. Continuous variables are reported as median (interquartile range). |  |  |  |  |
| P values indicate differences between not ICU and ICU admitted COVID 19 patients. |  |  |  |  |
| <sup>a</sup>Values presented as median and interquartile range |  |  |  |  |
| <sup>b</sup>Values presented as number and % of the column total |  |  |  |  |

**Table 3.** Demographic and Clinical characteristics associated with intubation.

|  | No Intubated<br>(n=42) | Intubated<br>(n=55) | OR for Intubation<br>(95% CI) | P value |
| --- | --- | --- | --- | --- |
| **Demographics** |  |  |  |  |
| Age, yr<sup>a</sup> | 61 (50.35-71.9) | 69.8 (63.9-78.5) | 1.05 (1.01-1.09) | 0.005 |
| Male n (%)<sup>b</sup> | 38 (52.8) | 16 (64) | 1.59 (0.63-4.19) | 0.3 |
| African-American<sup>b</sup> | 56 (77.8) | 19 (76) | 0.9 (0.31-2.81) | 0.8 |
| BMI<sup>a</sup> | 28.9 (24.4-34.8) | 29.7 (25-32.9) | 1.0 (0.94-1.06) | 0.9 |
| Smoking history<sup>b</sup> | 32 (44.5) | 13 (52) | 1.35 (0.54-3.4) | 0.5 |
| **Comorbidities** |  |  |  |  |
| HTN<sup>b</sup> | 53 (73.6) | 20 (80) | 1.43 (0.49-4.70) | 0.5 |
| DMII<sup>b</sup> | 36 (50) | 15 (60) | 1.5 (0.6-3.87) | 0.4 |
| COPD<sup>b</sup> | 8 (11.1) | 5 (20) | 2.0 (0.55-6.71) | 0.3 |
| CKD>2<sup>b</sup> | 25 (34.7) | 12 (48) | 1.73 (0.68-4.39) | 0.2 |
| ESRD<sup>b</sup> | 5 (6.9) | 3 (12) | 1.82 (0.35-8.07) | 0.4 |
| CAD<sup>b</sup> | 17 (23.6) | 9 (36) | 1.82 (0.66-4.82) | 0.2 |
| CVA<sup>b</sup> | 14 (19.4) | 7 (28) | 1.61 (0.54-4.53) | 0.4 |
| VTE<sup>b</sup> | 13 (18) | 3 (12) | 0.61 (0.13-2.14) | 0.5 |
| 2 or more<br>comorbidities<sup>b</sup> | 44 (61.1) | 18 (72) | 2.01 (0.74-6.08) | 0.2 |
| 3 or more<br>comorbidities<sup>b</sup> | 34 (47.2) | 16 (64) | 1.98 (0.79-5.25) | 0.2 |
| Definition of Abbreviations: HTN= Hypertension; DMII= Diabetes Mellitus Type II; COPD= Chronic Obstructive Pulmonary Disease; CKD>2= Chronic Kidney Disease; ESRD= End-Stage Renal Disease; CAD= Coronary Artery Disease; CVA= Cerebrovascular Accident; VTE= Venous Thromboembolism. Continuous variables are reported as median (interquartile range). |  |  |  |  |
| P values indicate differences between not intubated and intubated COVID 19 patients. |  |  |  |  |
| <sup>a</sup>Values presented as median and interquartile range |  |  |  |  |
| <sup>b</sup>Values presented as number and % of the column total |  |  |  |  |

## MAIN RESULTS

### COVID-19 patients with high circulating MT-DNA are at higher risk for mortality

To assess MT-DNA levels in COVID-19 patients we utilized a previously established in-situ quantitative PCR method (Scozzi et al., 2019) to measure the accumulation of fragments derived from the mitochondrial encoded gene cytochrome b (MT-CYTB) within cell-free circulating plasma. Plasma MT-CYTB levels were elevated in those subjects who died from COVID-19 [7.881 (6.897 - 8.488, n=25)] compared to those who survived [7.082 (6.650 - 7.693), n=72, p=0.009, **Figure 2A**] and were associated with an increased risk for mortality on univariate logistic regression (OR 1.960, 95% CI 1.236 - 3.28, p = 0.006). For MT-CYTB, area under the curve (AUC) for mortality was 0.68 (95% CI 0.54 – 0.81, **Figure 2B**). On multivariable logistic regression, plasma MT-CYTB levels remained an independent risk factor for mortality when adjusted for age, sex and 2 or more comorbidities (OR_adj_, 1.92, 95% CI 1.154 - 3.355, p = 0.015, **Table 4**).

**Table 4.** Multivariate analysis associated with mortality.

|  | **Adjusted OR for Mortality**<br>**(95% CI)** | **P value** |
| --- | --- | --- |
| MT-CYTB | 1.921 (1.154 - 3.355) | 0.015 |
| Age | 1.074 (1.026 - 1.131) | 0.003 |
| Sex, (male) | 1.380 (0.460-4.362) | 0.570 |
| 2 or more comorbidities | 2.578 (0.611-14.200) | 0.225 |
| N =97 (25 mortality events) |  |  |

**Figure 2.**
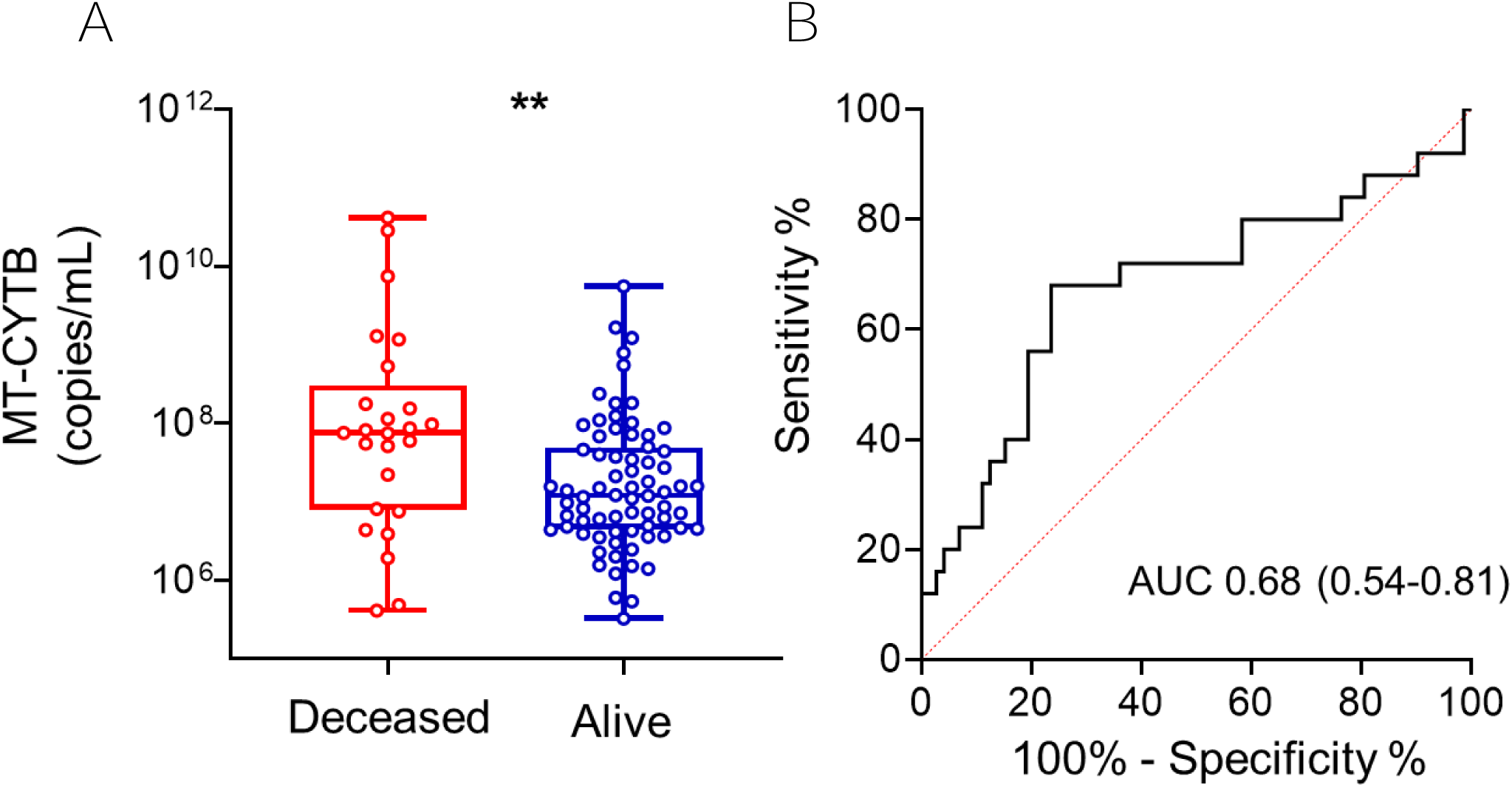
High Circulating MT-DNA levels predict a higher risk of mortality in COVID-19 patients. Plasma for determination of circulating levels of MT-CYTB was obtained at time of hospital presentation **(A)** Box and whiskers plot of MT-CYB levels in relation to mortality status in COVID-19 patients. **(B)** Receiver operating characteristic (ROC) curves in predicting the outcome mortality based on MT-CYTB levels. (\*\**P* < .01)

### COVID-19 patients with high circulating MT-DNA are likely to require ICU admission and intubation

Plasma MT-CYTB levels were elevated in those subjects with COVID-19 who required an ICU admission [7.779 (6.882 - 8.257), n=55] compared to those who were not admitted to the ICU [6.856 (6.544 - 7.293), p<0.0001, n=42, **Figure 3A**]. MT-CYTB levels were associated with ICU admission on univariate logistic regression (OR 3.335, 95% CI 1.890 - 6.564, p < 0.001). For MT-CYTB, AUC for ICU admission was 0.75 (95%CI 0.65 – 0.85, **Figure 3B**). On multivariable logistic regression, plasma MT-CYTB levels remained an independent risk factor for ICU admission after adjusting for age, sex and 2 or more comorbidities (OR_adj_, 3.146, 95% CI 1.652 - 6.972, p = 0.002, **Table 5**). Notably, we also made similar findings for another mitochondrial DNA encoded gene MT-COX3 (**Figure S1D and E**, OR_adj_ for age, sex and 2 or more comorbidities, 1.561, 95% CI 1.170 - 2.188, p = 0.005, AUC of COX3 for ICU admission 0.69, 95% CI 0.58 - 0.79).

**Table 5.** Multivariate analysis associated with ICU admission.

|  | **Adjusted OR for Mortality**<br>**(95% CI)** | **P value** |
| --- | --- | --- |
| MT-CYTB | 3.146 (1.652 - 6.972) | 0.002 |
| Age | 1.113 (1.062 - 1.180) | <0.0001 |
| Sex, (male) | 2.060 (0.688 - 6.478) | 0.201 |
| 2 or more comorbidities | 0.465 (0.113 – 1.713) | 0.264 |
| N =97 (55 ICU events) |  |  |

**Figure 3.**
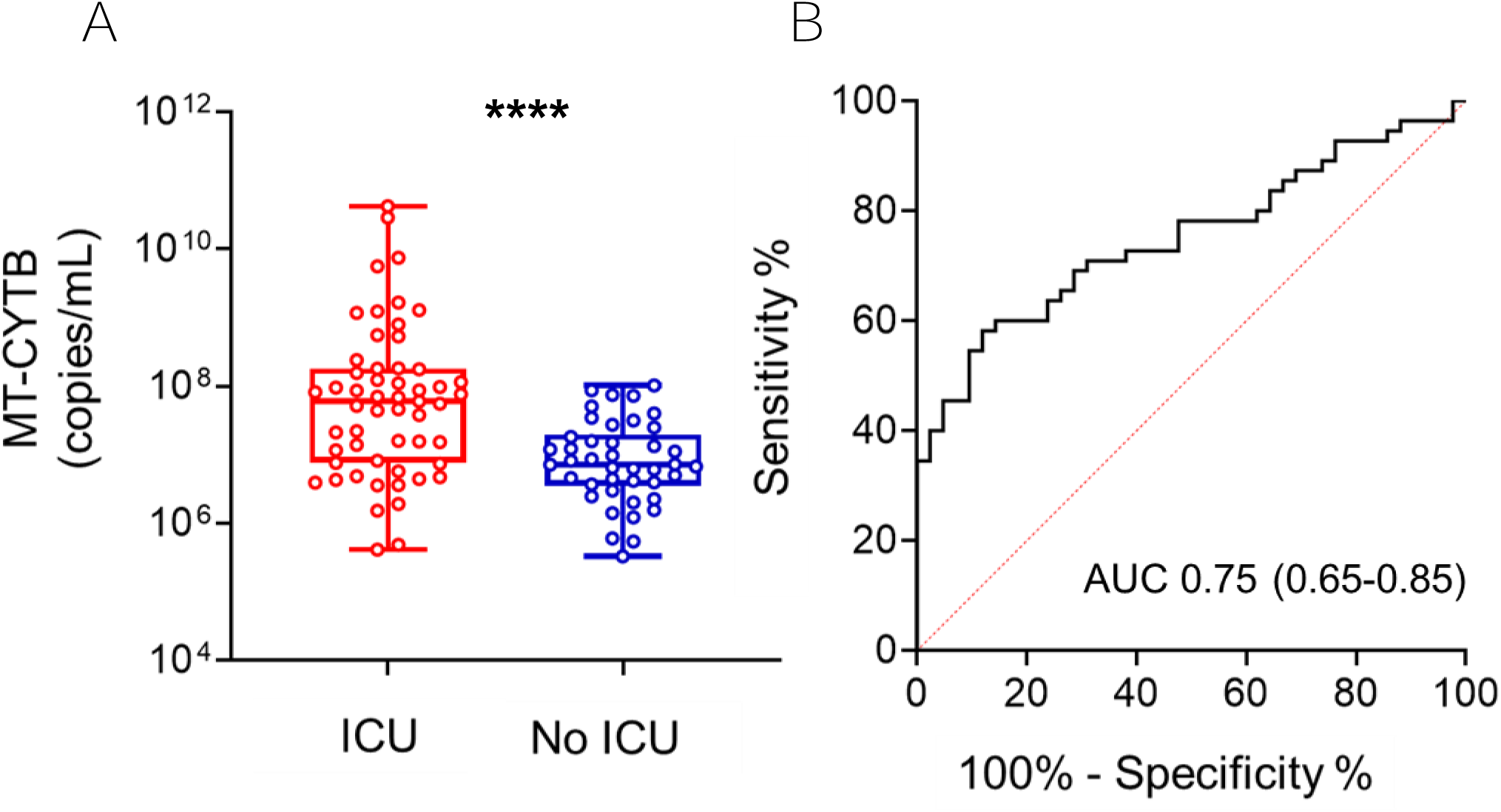
High circulating MT-DNA levels predict a higher risk of ICU requirement in COVID-19 patients. Plasma for determination of circulating levels of MT-CYTB was obtained at time of hospital presentation (**A)** Box and whiskers plot of MT-CYB levels in relation to ICU admission in COVID-19 patients. **(B)** Receiver operating characteristic (ROC) curves in predicting the outcome ICU based on MT-CYTB levels. (\*\*\*\**P* < .0001).

Similarly, plasma MT-CYTB levels were elevated in those subjects with COVID-19 who required intubation [8.192 (7.830 - 9.101, n=25)] versus those who did not require intubation [6.963 (6.603 - 7.597), n=72, p<0.0001, **Figure 4A**]. Plasma MT-CYTB levels were associated with intubation on univariate logistic regression (OR 6.251, 95% CI 3.010 - 16.110, p < 0.0001). For MT-CYTB, AUC for intubation was 0.86 (95%CI 0.76 – 0.95, **Figure 4B**). On multivariable logistic regression, plasma MT-CYTB levels remained an independent risk factor for intubation (OR_adj_, 5.881, 95% CI 2.814 - 15.520, p < 0.0001, **Table 6**). Similar findings were observed with MT-COX3 [**Figure S1F and S1G**, OR_adj_ for age, sex and 2 or more comorbidities, 2.759, 95% CI 1.791 - 4.921, p < 0.0001, AUC for intubation 0.80 (95% CI 0.69 - 0.91)].

**Table 6.** Multivariate analysis associated with intubation.

|  | **Adjusted OR for Mortality**<br>**(95% CI)** | **P value** |
| --- | --- | --- |
| MT-CYTB | 5.881 (2.814 - 15.52) | <0.0001 |
| Age | 1.059 (1.008 - 1.119) | 0.03 |
| Sex, (male) | 1.339 (0.381 - 4.920) | 0.649 |
| 2 or more comorbidities | 0.831 (0.166 - 4.368) | 0.821 |
| N =97 (25 intubation events) |  |  |

**Figure 4.**
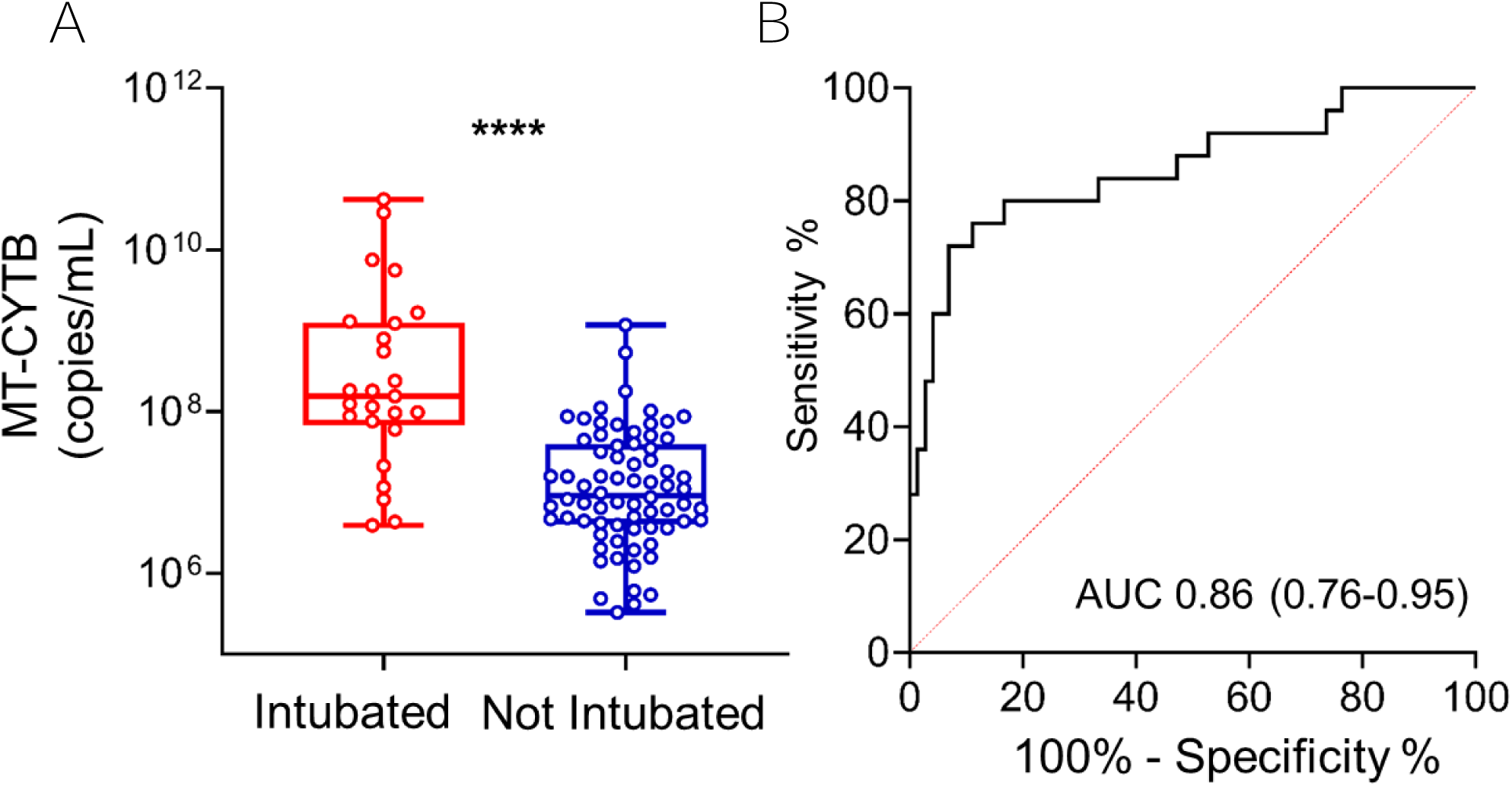
High circulating MT-DNA levels predict a higher risk of intubation in COVID-19 patients. Plasma for determination of circulating levels of MT-CYTB was obtained at time of hospital presentation **(A)** Box and whiskers plot of MT-CYB levels in relation to requirement for intubation in COVID-19 patients. **(B)** Receiver operating characteristic (ROC) curves in predicting the outcome intubation based on MT-CYTB levels. (\*\*\*\**P* < .0001).

### Circulating MT-DNA levels show similar or improved accuracy over clinically established measurements of inflammation typically used in COVID-19 patients

Plasma MT-CYTB levels had a similar AUC for mortality when compared to LDH, ferritin or D-dimer levels, and were better than CRP drawn within the first 24 hours of presentation (**Figure 5A**). Importantly, the AUC for plasma MT-CYTB levels was superior to CRP, LDH, ferritin and D-dimer levels when predicting the need for an ICU admission (**Figure 5B**) or intubation (**Figure 5C**). A similar pattern was identified for MT-COX3 when compared to CRP, LDH, ferritin and D-dimer levels for predicting the need for an ICU admission (**Figure S2B**) but was superior to these clinically utilized markers for the need for intubation (**Figure S2C**).

**Figure 5.**
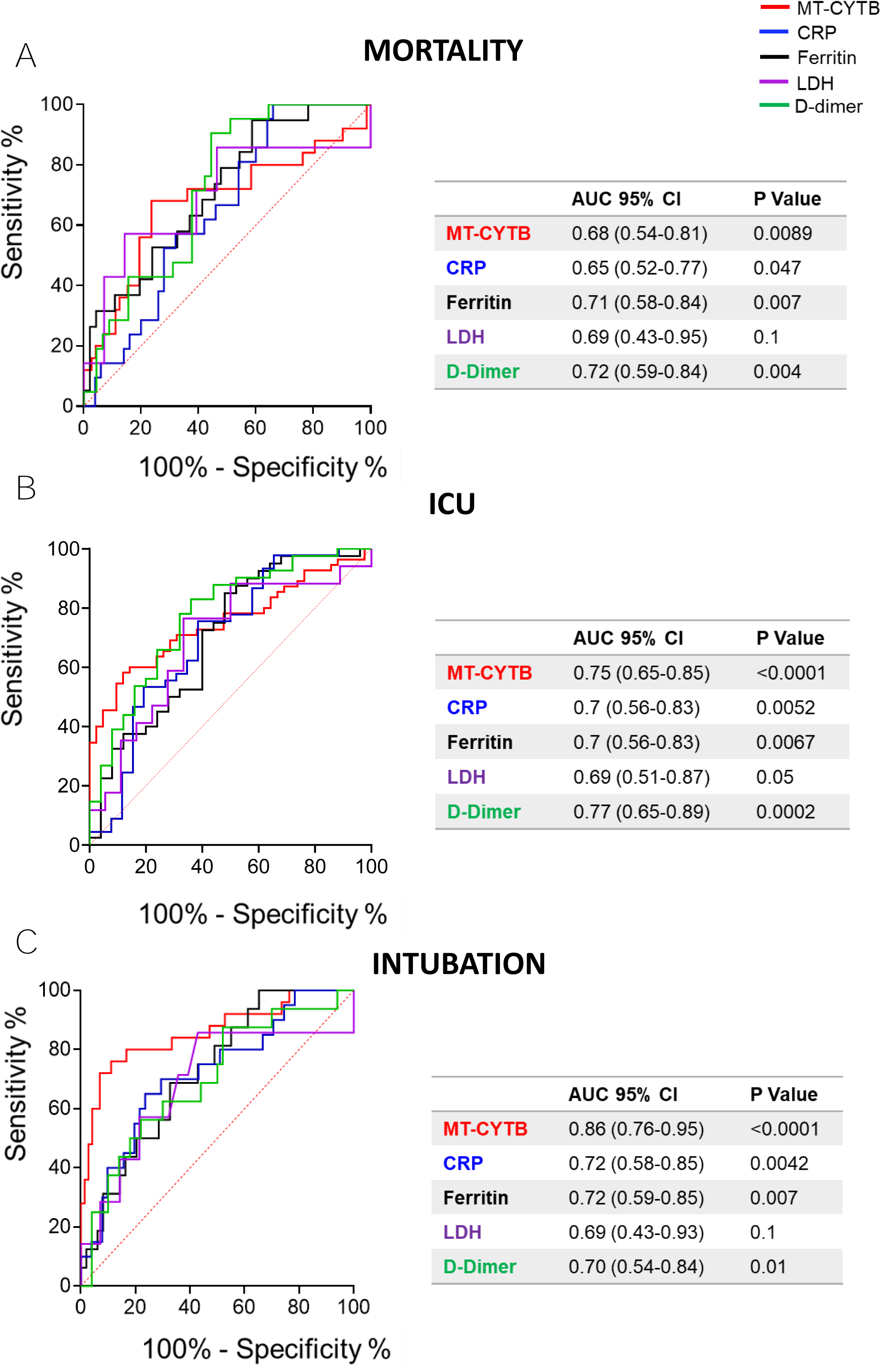
Circulating MT-DNA levels have a similar or improved accuracy over clinically utilized biomarkers for outcomes of severity in COVID-19. Blood samples for determination of biomarkers levels were collected within 24 hours from hospital presentation. Receiver operating characteristic (ROC) curves in predicting the outcome **(A)** mortality, **(B)** admission to ICU and **(C)** Intubation based on MT-CYTB (red), reactive C protein (CRP) (blue), Ferritin (black), Lactic acid dehydrogenase (LDH) (purple) and D-Dimer (green) levels. Area under the curve (AUC) with 95% CI and P values for the different biomarkers are summarized in the corresponding tables.

### Circulating MT-DNA levels correlate with other emerging markers of COVID-19 severity

Plasma MT-CYTB levels moderately correlated with concurrently measured levels of IL-6, which has been implicated in the pathogenesis of COVID-19 [r=0.389, 95% CI 0.194 – 0.554, p = 0.0001, n=92, **Figure 6A**]. Similarly, MT-CYTB levels highly correlated with plasma sC5b-9, which is a marker of complement activation and suggests the formation of a membrane attack complex (MAC) [r=0.49, 95% CI 0.31 – 0.63, p < 0.0001, n=95, **Figure 6B**]. MT-CYTB also correlated with the neutrophil-to-lymphocyte ratio [r=0.37, 95% CI 0.17 – 0.54, p = 0.0003, n=90, **Figure 6C]**. Plasma MT-CYTB levels had a similar or/improved accuracy compared to IL-6 for mortality (**Figure S3A**), ICU admission (**Figure S3B**) and intubation (**Figure S3C**).

**Figure 6.**
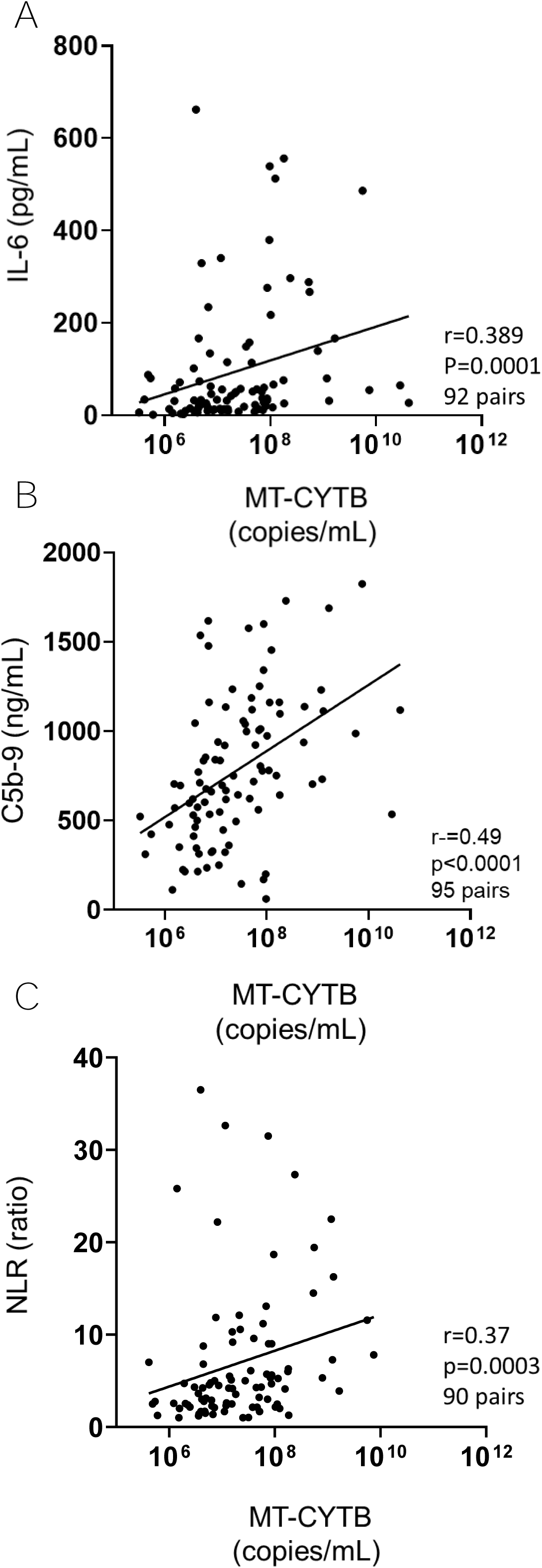
Circulating MT-DNA levels correlate with other emerging markers of inflammation found in COVID-19 patients. Blood samples for determination of inflammatory indicators were collected within 24 hours from hospital presentation. Scatter plots showing the correlation between MT-CYTB and **(A)** IL-6, **(B)** C5b-9 (terminal complement complex and **(C)** Neutrophil to Lymphocyte ratio (NLR).

We also observed significant correlations with other biomarkers that have been implicated in the pathogenesis of COVID-19 such as MIG [r=0.31, 95% CI 0.11 – 0.49, p = 0.002, n=93, **Figure S4A**], MCP-1 [r=0.25, 95% CI 0.04 – 0.44, p = 0.01, n=93, **Figure S4B**], IP-10 [r=0.25, 95% CI 0.05 – 0.44, p = 0.01, n=94, **Figure S4C**], IL-1RA [r=0.43, 95% CI 0.24 – 0.59, p < 0.0001, n=94, **Figure S4D**] and IL-2R [r=0.28, 95% CI 0.08 – 0.46, p = 0.005, n=94, **Figure S4E**]. Similarly, the levels of HGF highly correlated with MT-CYTB in COVID-19 [r=0.47, 95% CI 0.29 –0.62, p < 0.0001, n=94, **Figure S4F**].

## DISCUSSION

Here we observe that high levels of circulating MT-DNA are an early independent risk factor for severe illness and mortality in hospitalized patients with COVID-19, after adjusting for age, sex and comorbidities. Importantly, we made similar findings for two MT-DNA genes, MT-CYTB and MT-COX3, although the latter target did not reach significance for mortality risk. The reasons for this incongruence are not clear. However, when compared to MT-CYTB, we generally detected lower copy numbers for MT-COX3, suggesting the possibility that this part of mitochondria genome is more intrinsically unstable when exposed to plasma (Scozzi et al., 2019).

Although our studies were not specifically designed to identify the mechanisms that drive peripheral blood MT-DNA accumulation, significant correlations between LDH and IL-6 with MT-DNA levels point to a potential deleterious role for cellular necrosis in COVID-19 pathophysiology. LDH release and IL-6 production are reported indicators of cellular necrosis (Chan et al., 2013; Vanden Berghe et al., 2006). A specific form of necrosis, necroptosis, has been demonstrated to induce the release of damaged mitochondria (Maeda and Fadeel, 2014). Necroptosis is characterized by the eventual loss of plasma membrane integrity due to the kinase activity of the receptor-interacting proteins (RIPK)1 and RIPK3 (He et al., 2009) and has been reported to act as a host defense mechanism when apoptotic death pathways are disabled by viral infection (Cho et al., 2009). For example, in a mouse model of influenza infection, necroptosis was shown to inhibit viral replication but with deleterious consequences to bronchial epithelial integrity (Rodrigue-Gervais et al., 2014). In humans, both H5N1 and H1N1 influenza-induced acute respiratory distress syndrome have been shown to be associated with necrotic cell death within the distal pulmonary epithelia (Gu and Korteweg, 2007; Mauad et al., 2010). Intriguingly, the accessory protein open reading frame 3a (Orf3a) expressed within the highly related SARS-CoV-1 genome has been recently demonstrated to induce necroptosis through activating RIPK3 (Yue et al., 2018). SARS-CoV-2 also expresses an Orf3a accessory protein (Bojkova et al., 2020). However, whether SARS-CoV-2 Orf3a promotes necroptosis has yet to be determined. Some MT-DNA release may also result from innate immune cell activation. For example, human neutrophils following activation extrude their MT-DNA due to an inability to complete mitophagy (Caielli et al., 2016). Additionally, neutrophil extracellular traps (NETs), which have been found in the lungs of COVID-19 patients (Zuo et al., 2020), are rich in MT-DNA (Yousefi et al., 2009). Neutrophils have also been identified as undergoing metabolic reprogramming in the context of SARS-CoV-2 (McElvaney et al., 2020). In addition to neutrophils, activated platelets also release mitochondria, which in turn can induce NET generation (Caudrillier et al., 2012), possibly contributing to pulmonary pro-thrombotic complications observed in patients with severe COVID-19 illness (Middleton et al., 2020). The source and mechanisms of MT-DNA release in response to SARS-CoV-2 infection is a subject of future investigation.

Multiple clinically established biomarkers such as LDH, Ferritin, CRP and D-dimers are currently being evaluated to assess the risk of clinical deterioration from a COVID-19 diagnosis (Cummings et al., 2020; Luo et al., 2020; McElvaney et al., 2020; Messner et al., 2020; Song et al., 2020; Wu et al., 2020). However, as products of gene expression in response to both acute and chronic stimuli they tend to be largely non-specific measures of systemic inflammation with the exception of LDH, a marker of cell death. Nevertheless, LDH can also be released by cells undergoing apoptosis, a predominantly anti-inflammatory form of cell death (Kumar et al., 2018). In contrast, there are accumulating observations that high amounts of MT-DNA release are specifically generated by necrotic cells (Vringer and Tait, 2019). Additionally, high MT-DNA levels have been shown to be associated with acute lung injury, in multiple, independent cohorts (Faust et al., 2020; Huang et al., 2020; Nakahira et al., 2013). Here, we observed that MT-DNA levels are approximately 10-fold higher in COVID-19 patients who developed severe pulmonary dysfunction or eventually died suggesting it is at least as sensitive a biomarker as other clinically established and exploratory indicators used in prediction models. To conduct plasma MT-DNA measurements we employed a rapid PCR assay technique that takes about 60 minutes to complete due to the elimination of the DNA purification step. In resource-limited settings, this can be especially important given the current necessity to identify subjects at a higher risk of clinical deterioration. Additionally, PCR-based assays measuring MT-DNA tend to be less cost prohibitive and do not need specialized equipment, facilitating easy implementation. Nevertheless, whether MT-DNA levels are equivalent or better measures than routinely obtained laboratory measurements will require additional validation with independent cohorts.

Based on our data from this study, it is not possible to clearly determine if circulating MT-DNA contributes to the pathogenesis of COVID-19 disease. Nevertheless, cell-free MT-DNA is itself a signatory marker for the release of other MT DAMPs, which collectively drive pro-inflammatory cytokine expression through the engagement of PRRs on innate immune cells (Kulkarni et al., 2020a). MT DAMPs drive IL-6 expression by macrophages (Maeda and Fadeel, 2014) and stimulate IL-8 release by neutrophils (Hauser et al., 2010). Notably, IL-6 suppresses lymphopoiesis (Maeda et al., 2005) while IL-8 promotes neutrophil release from the bone marrow (Terashima et al., 1998) and therefore, could possibly explain our observed correlation between MT-DNA levels and elevations in the neutrophil to lymphocyte ratio. We additionally noted a significant correlation between membrane attack complex and MT-DNA levels. Extracellular mitochondria have been reported to activate complement. Mannan-binding lectin has been observed to bind to cell free mitochondria resulting in C3 consumption in the peripheral blood of mice (Brinkmann et al., 2013). Therefore, it is interesting to note that C3 consumption is a general sign of C3 convertase activity, which would be a prerequisite for the downstream generation of membrane attack components (Leslie and Nielsen, 2004). Along those lines, complement activation has been implicated in the pathogenesis of COVID-19-related end-organ damage, including acute lung injury (Java et al., 2020; Magro et al., 2020), with potential therapeutic implications (Kulasekararaj et al., 2020; Smith et al., 2020). Finally, MT DAMPs, unlike other inflammatory markers linked to poor outcomes of COVID-19 afflicted patients, have been reported to directly cause acute pulmonary dysfunction and tissue damage (Lee et al., 2017). Administration of MT DAMPs into the blood stream or pulmonary airways of rodents promotes acute lung injury mediated by neutrophil chemotaxis and reactive oxygen species generation to mitochondrial formylated peptides (Hauser et al., 2010; Scozzi et al., 2019). In these studies, the formylated peptide receptor-1 inhibitor cyclosporine H was shown to inhibit MT DAMP-mediated acute lung injury. Given ongoing clinical trials that target the effects of IL-6, complement activation and NETs in COVID-19 patients (Desilles et al., 2020; Kulkarni et al., 2020b; Maes et al., 2020; Rilinger et al., 2020; Smith et al., 2020) future approaches that block necroptosis or MT DAMP recognition could also be warranted.

Our study has several limitations. First, we were unable to concurrently enroll a COVID-19 negative group with severe respiratory disease. Our center, as many others experienced a decline in hospital presentations for illnesses other than COVID-19 during this time making a concurrent, adequately matched control group not possible. Additionally, as our samples were drawn as a part of a prospective study of subjects with known SARS-CoV-2 positivity, any other adequately matched control group would not be entirely comparable, as it was not concurrently drawn. Importantly, we do not aim to propose that MT-DNA levels are different in those with COVID-19 compared to those with acute lung injury due to other etiologies. Multiple emerging studies suggest that biomarkers may be no different in subjects with COVID-19-induced lung injury versus those due to other etiologies (Sinha et al., 2020b). However, the strength of our study lies in an assay that can potentially identify subjects presenting with COVID-19 who are likely to develop adverse outcomes during the course of their hospitalization, which becomes increasingly important when resources are constrained as has been unfortunately common during this pandemic (Barrett et al., 2020; Litton et al., 2020; Moghadas et al., 2020). Second, this study does not assess how MT-DNA correlates with viral load in COVID-19, which is especially important in the context of deciding when to intervene with targeted therapeutics (Catanzaro et al., 2020; Du and Yuan, 2020; Maggi et al., 2020). Therefore, further studies will be necessary to evaluate whether it may be a predictor of response to treatment (Sinha et al., 2020a). Third, decision-making in the ICU changed over the course of our enrollment. For example, there was a tendency towards intubating earlier in the course of the critical illness in the initial months of the COVID-19 pandemic. However, we identified MT-DNA as a predictor of severity utilizing mortality as a primary endpoint, and then ICU admission and the requirement for intubation as independent, secondary endpoints.

In summary, we demonstrate that MT-DNA measured early in the disease course can predict survival status, the requirement for intensive level care, and the need for endotracheal intubation. We also show that MT-DNA levels are associated with exploratory biomarkers implicated in the pathogenesis of COVID-19-related morbidity and mortality. Further studies will be needed to discern the contribution of MT DAMPs such as MT-DNA to COVID-19 pathogenesis, as well as to understand whether MT-DNA and other inflammatory mediators, such as activated complement, act synergistically to promote cellular injury.

## ACKNOWLEDGMENTS

A.E.G. is supported by Washington University Institute of Clinical Translational Sciences (ICTS) COVID-19 Research Program, The Barnes Jewish Foundation, NIH R01HL094601 and NIH P01AI116501. H.S.H is supported by NIH K08HL148510 and the Children’s Discovery Institute. J.A.O., C.G. and P.A.M. is supported by Washington University ICTS grant UL1TR002345 from the National Center for Advancing Translational Sciences (NCATS) of the NIH. The content is solely the responsibility of the authors and does not necessarily represent the official view of the NIH. Sample procurement and patient outcome data collection was supported by the Washington University ICTS NIH grant UL1TR002345.

## AUTHOR CONTRIBUTIONS

Conceptualization: H.S.K., A.E.G., M.I., Methodology: D.S., M.C., C.G., H.S.K., A.E.G., Software: D.S., M.C., H.S.K., Validation: J.O., A.A.R., C.G., Formal Analysis: D.S., M.C., V.P., M.R., A.R., R.A., L.A., H.S.K., A.E.G., Investigation: L.M., D.Z., J.Z., Data Curation: L.M., D.S., M.C., H.S.K., A.E.G., Writing – Original Draft: D.S., M.C., H.S.K., A.E.G., Writing – Review & Editing: D.S., M.C., L.M., J.H., C.G., A.R., Z.L., R.H., M.I., D.K. H.S.K., A.E.G., Visualization: D.S., M.C., Supervision: H.S.K., A.E.G., Project Administration: H.S.K., A.E.G., Funding Acquisition: H.S.K., R.R.H., D.K., A.E.G.

## METHODS

### Study Design, Settings and Participants

This prospective cohort study utilized cell-free plasma samples that had been prospectively collected from 97 adult patients with confirmed COVID-19 presenting to the Barnes-Jewish Hospital from March 15, 2020, to April 24, 2020. Diagnosis of COVID-19 was based on a positive nasopharyngeal swab test.

### Study Approval

The study was approved by the Washington University School of Medicine Institutional Review Board (ID#202004091 and #202003085). Written informed consent was obtained from all subjects.

### Outcome Definition

Subjects were followed through June 29, 2020. The primary outcome was mortality. The secondary outcomes included (a) the need for an ICU admission and, (b) endotracheal intubation. These outcomes were abstracted utilizing an honest broker system from electronic medical records.

### Sample Collection and Processing

MT-DNA and other cytokines were measured in cell-free plasma of subjects within the first 24 hours of emergency department presentation. Blood samples were collected in EDTA-containing vacutainers (BD Biosciences, San Jose, CA) and subjected to 2 rounds of centrifugation to generate platelet poor plasma. First at 2,500 x g for 20 mins to generate plasma. The plasma was removed from vacutainers and centrifuged at 13,000 ⨯ g in sterile nuclease-free eppendorf tubes (ThermoFisher Scientific, Waltham, MA) for 10 min to remove platelets. These platelet poor specimens were then immediately stored at −80°C until further analysis. Concurrently measured clinical markers (I.e., C-reactive protein, ferritin, lactate dehydrogenase, D-dimer) were obtained from the electronic medical record.

### MT-DNA quantification

MT-DNA quantification Real-Time PCR was performed in a BioRad CFX-Connect machine using reaction mixture containing 0.1 μL of cell-free plasma, 10 μL iQ SYBR Green Supermix (Bio-Rad), 0.5 μL of 5μM forward and reverse primers and 8.9 μL H2O. Assays were performed in triplicate under the following conditions: 1 cycle at 95°C for 3 min, then up to 40 cycles at 95°C for 10 sec and 55°C for 30 sec and then a melt curve was performed from 65°C to 95 °C (0.5 °C every 5 sec). Primers for Human cytochrome B (CYB; forward 5_′_-ATGACCCCAATACGCAAAAT-3_′_ and reverse 5_′_-CGAAGTTTCATCATGCGGAG-3_′_), Human cytochrome C oxidase subunit III (COX3: forward 5_′_-ATGACCCACCAATCACATGC-3_′_ and reverse 5_′_-ATCACATGGCTAGGCCGGAG-3_′_) were synthesized by Integrated DNA technologies (IDT, Coralville, IA). Copy number was estimated by comparison to a real-time PCR standard amplification curve generated from known amounts of purified human MT-DNA and analyzed after a log10 transformation.

### Quantification of cytokines and complement activation

Cell-free plasma was analyzed using a Cytokine 35-Plex Human Panel, which provide simultaneous measurement of 35 cytokines (ThermoFisher Scientific, Waltham, MA). The assay was performed in accordance with manufacturer’s instructions with each subject sample performed in duplicate and then analyzed on a Luminex FLEXMAP 3D instrument. Complement activation was assessed in cell-free plasma specimens (not previously thawed) using the soluble C5b-9 (sC5b-9) assay (BD OptEIA Human C5b-9 ELISA set, Franklin Lakes, NJ, USA).

### Statistical Analysis

Continuous variables were reported as median (interquartile range). The predictive value of each biomarker was expressed as area under the curve (AUC) derived from the relative ROC curve. Spearman r coefficient was calculated to estimate the correlation between two continuous variables. Univariate logistic regression analysis was used to calculate the unadjusted odds ratios. Independent variables known to have a biological relationship with the outcomes were selected a priori based on existing literature for the multivariate logistic model to calculate the adjusted odds ratios (Lederer et al., 2019; Petrilli et al., 2020; Williamson et al., 2020). Model diagnostics were performed with Corrected Akaike Information Criterion (AICc), log-likelihood, and model deviance. All the statistical analyses were calculated, and all figures were prepared using Graphpad Prism 8.4.2 (GraphPad Software Inc., La Jolla, CA). For all the analyses, a p value <0.05 was considered significant.

## SUPPLEMENTAL INFORMATION

**Suppl. Fig. 1.**
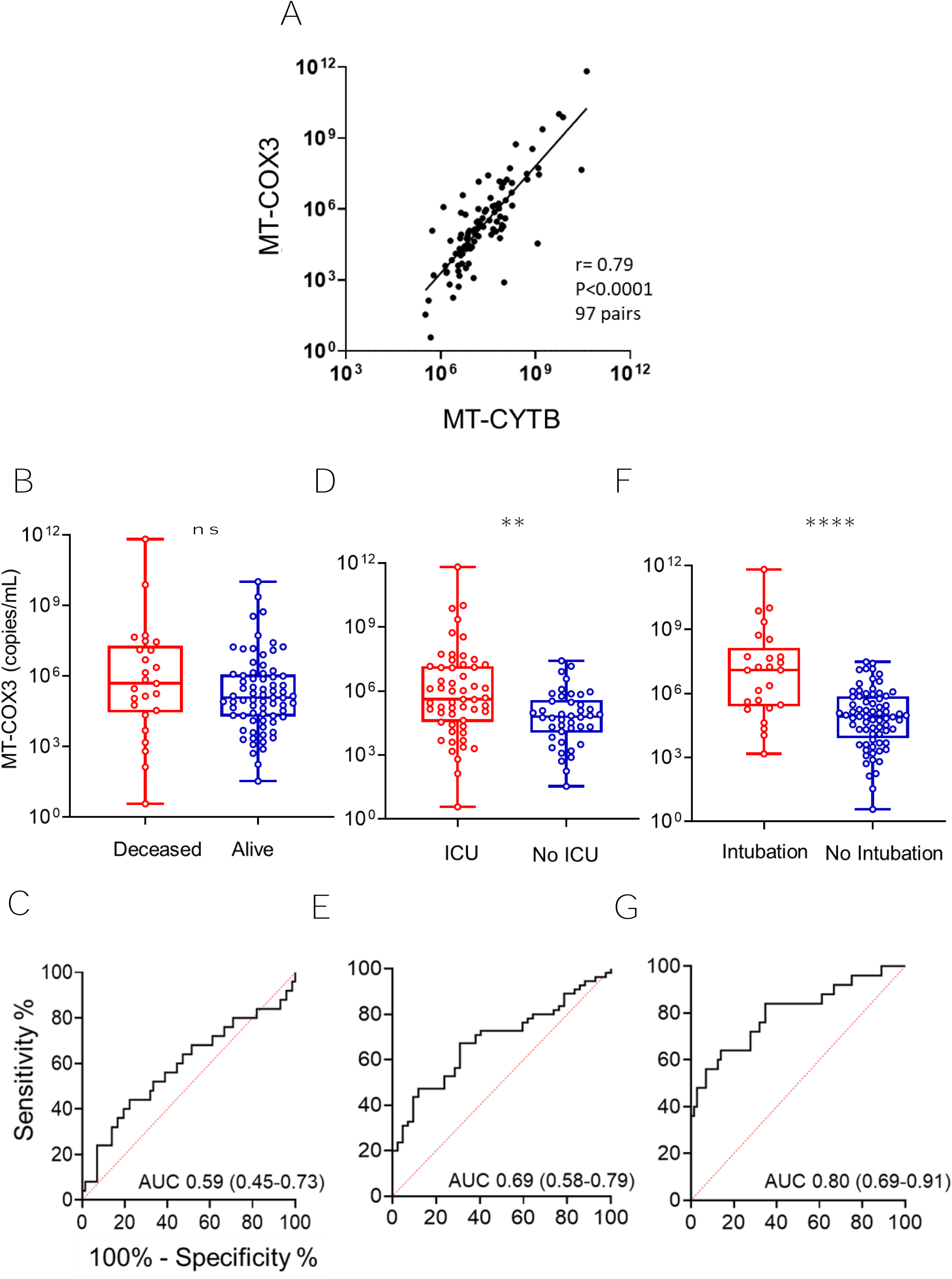
Comparable trends observed using MT-COX3 as an independent measure of MT-DNA. **(A)** Scatter plots showing the correlation between MT-CYTB and MT-COX3 measurements. Box and whiskers plot of MT-COX3 levels in relation to **(B)** mortality status, **(D)** ICU admission and **(F)** intubation in COVID-19 patients. Receiver operating characteristic (ROC) curves in predicting the outcome **(C)** mortality, **(E)** ICU admission and **(G)** intubation based on MT-COX3 levels in COVID-19. (\*\**P*□< □.01, \*\*\*\**P*□< □.0001)

**Suppl. Fig. 2.**
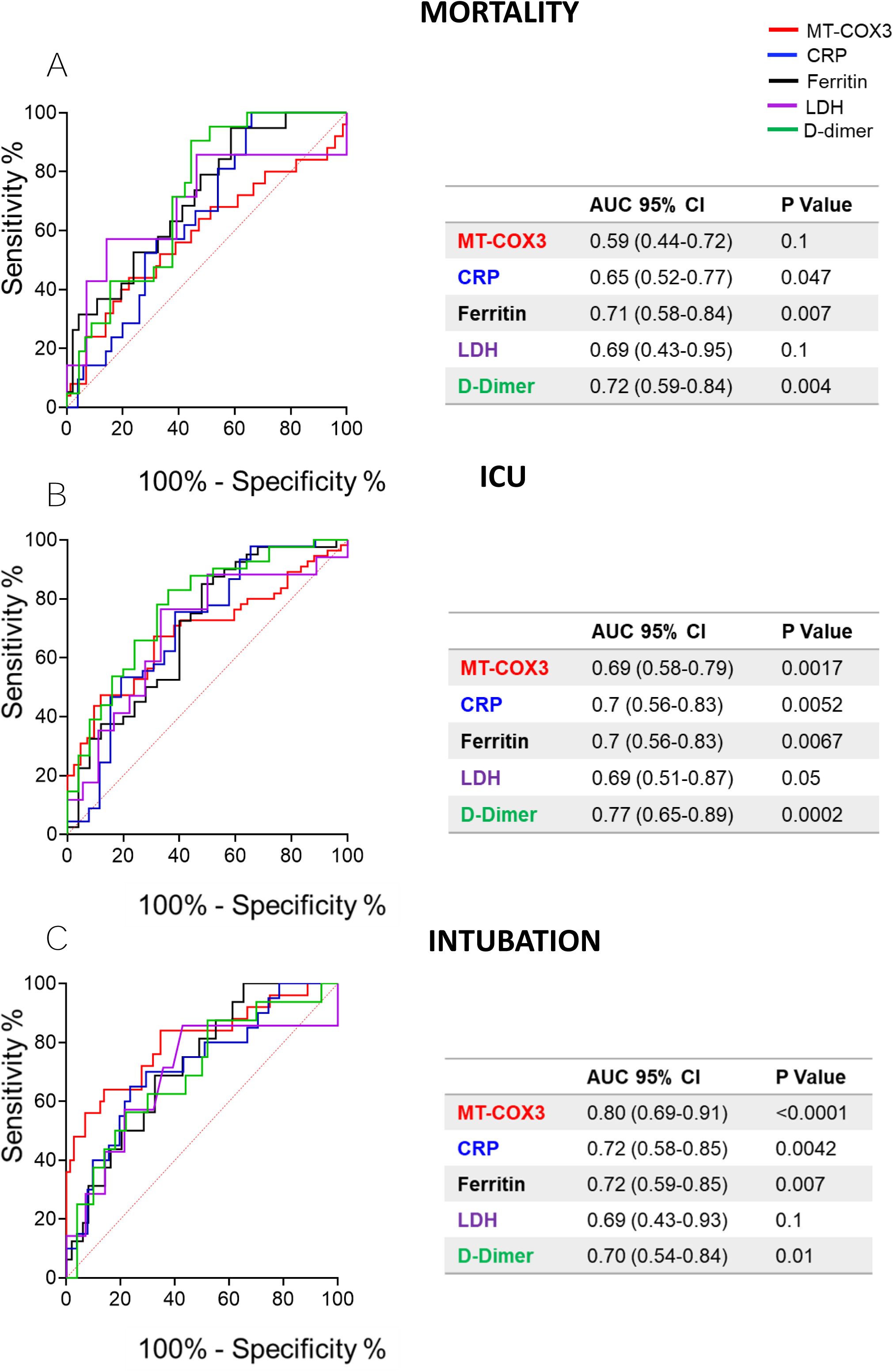
Comparison of the predictive values of MT-COX3 with current clinically used biomarkers of disease severity in COVID-19 patients. Blood samples for determination of biomarkers levels were collected within 24 hours from hospital presentation. Receiver operating characteristic (ROC) curves in predicting the outcome **(A)** mortality, **(B)** admission to ICU and **(C)** Intubation based on MT-COX3 (red), reactive C protein (CRP) (blue), Ferritin (black), Lactic acid dehydrogenase (LDH) (purple) and D-Dimer (green) levels. Area under the curve (AUC) with 95% CI and P values for the different biomarkers are summarized in the corresponding tables.

**Suppl. Fig. 3.**
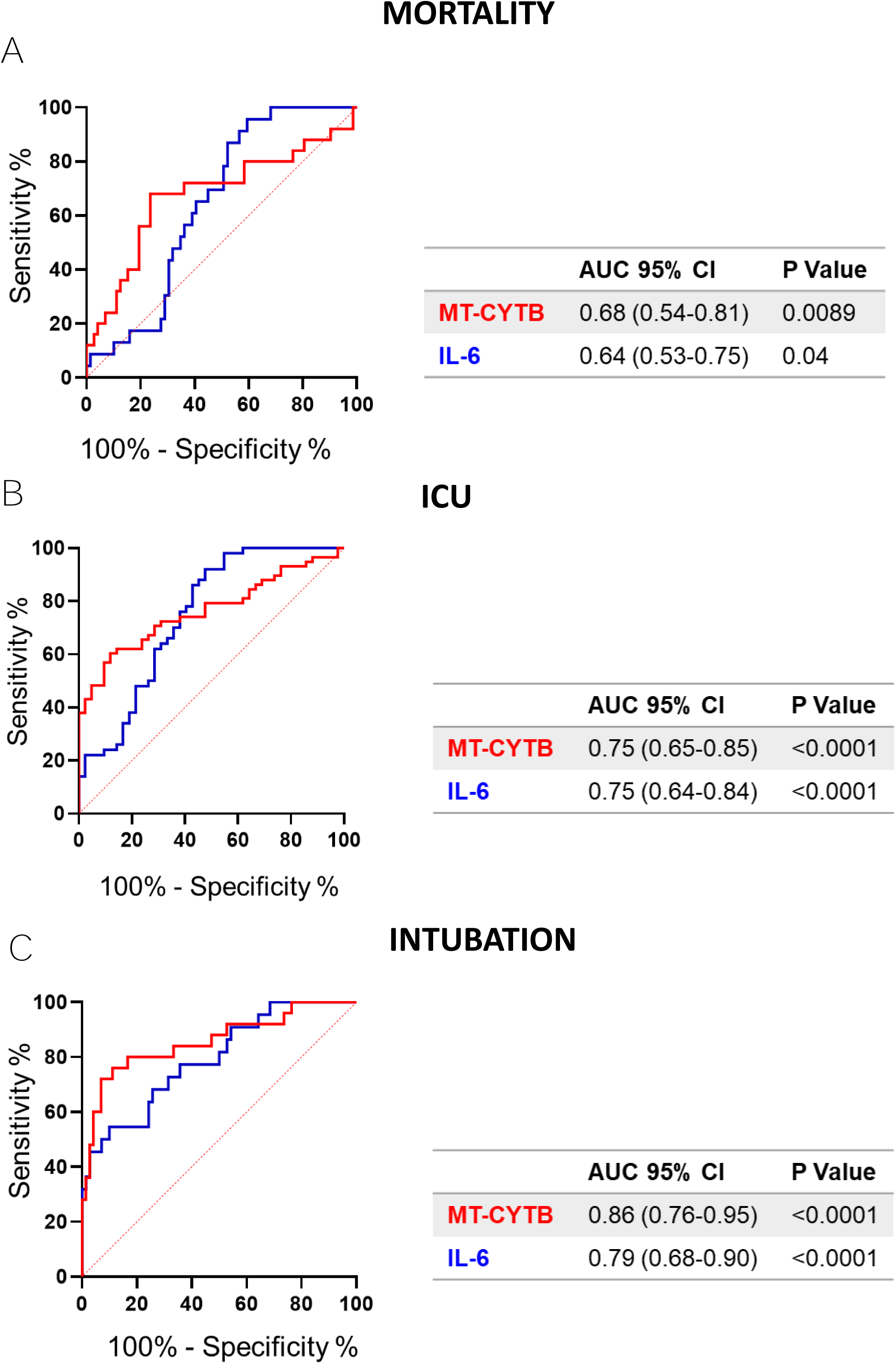
Comparison between the predictive values of MT-CYTB and IL-6 over outcomes of disease severity in COVID-19 patients. Receiver operating characteristic (ROC) curves in predicting the outcome **(A)** mortality, **(B)** admission to ICU and **(C)** Intubation based on MT-CYTB (red) and IL-6 (blue). Area under the curve (AUC) with 95% CI and P values are summarized in the corresponding tables.

**Suppl. Fig. 4.**
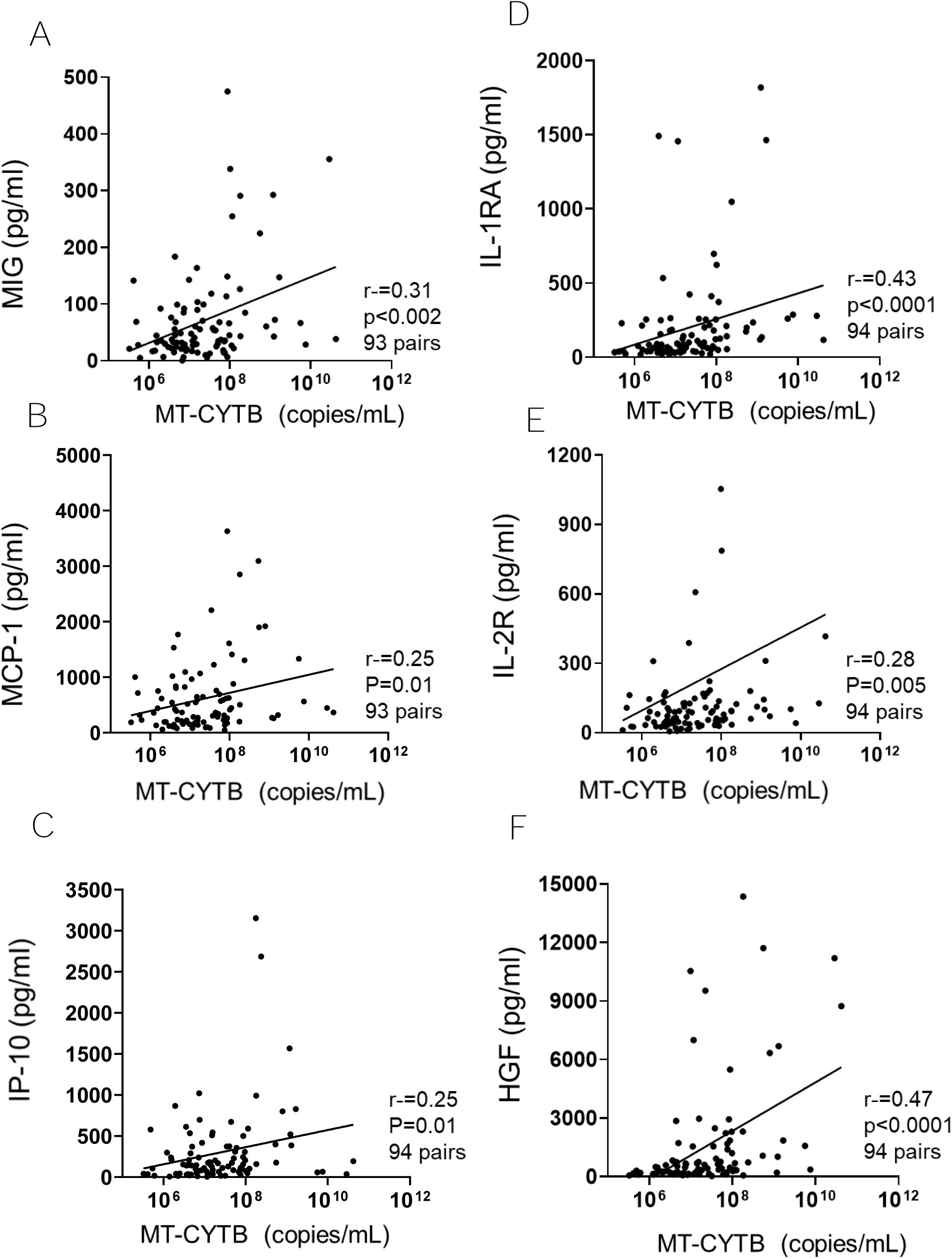
MT-CYTB levels positively correlate with emerging markers of disease severity in COVID-19 patients. Blood samples for determination of inflammatory indicators were collected within 24 hours from hospital presentation. Scatter plots showing the correlation between MT-CYTB and **(A)** MIG **(B)** MCP-1 **(C)** IP-10 **(D)** IL-1RA, **(E)** IL2R and **(F)** HGF.

